# GlycoAvatars: Bead-Coated Membrane Models for Studying the Cancer-Immune Cells Interactome

**DOI:** 10.1101/2025.01.15.633146

**Authors:** Andreia Miranda, Marta Relvas-Santos, Camila Lourenço, Eduardo Ferreira, Diogo M. Cunha, Carlos Palmeira, Lúcio Lara Santos, Pieta K Mattila, José Alexandre Ferreira

**Affiliations:** Experimental Pathology and Therapeutics Group, Research Center of IPO-Porto (CI-IPOP)/RISE@CI-IPOP (Health Research Network), Portuguese Oncology Institute of Porto (IPO-Porto)/Porto Comprehensive Cancer Center Raquel Seruca (Porto.CCC Raquel Seruca), Porto, Portugal; School of Medicine and Biomedical Sciences Abel Salazar (ICBAS), University of Porto, Porto, Portugal; Institute of Biomedicine, and MediCity Research Laboratories, University of Turku, Turku, Finland; Turku Bioscience, University of Turku and Åbo Akademi University, Turku, Finland; InFLAMES Research Flagship, University of Turku, Turku, Finland; Health School of University Fernando Pessoa, Porto, Portugal; Department of Immunology, Portuguese Oncology Institute of Porto (IPO-Porto), Porto, Portugal; Department of Surgical Oncology, Portuguese Oncology Institute of Porto (IPO-Porto), Porto, Portugal; GlycoMatters Biotech, Espinho, Portugal

**Keywords:** cancer glycosylation, glycoproteomics, interactomics, immune receptors, systems biology

## Abstract

Immature *O*-glycosylation of plasma membrane proteins, characterized by the presence of glycosites with a single GalNAc residue (the Tn antigen) rather than more extended and complex glycans, is a prominent feature of many advanced solid tumors across different origins and aetiologies. This form of glycosylation has been strongly associated with poor prognosis and is known to play a crucial role in the progression of the disease. Notably, it has been directly implicated in the crosstalk with the immune system, inducing tolerogenic phenotypes in both innate and adaptive immune cells. Despite this, the specific receptors involved in these interactions remain largely unidentified. To address this gap, we propose a novel high-throughput proteomics assisted strategy for characterizing the glycan interactome using plasma membrane-derived glycoprotein coated magnetic beads. Glycoproteins were isolated from plasma membranes of glycoengineered cell models with immature glycosylation and were then cloaked on the surface of magnetic beads, creating simplified cell “GlycoAvatars.” These GlycoAvatars were primarily employed to characterize the interactome between gastric and colorectal cancer cell lines and immune cells (dendritic cells and macrophages), providing a tool for identifying potentially relevant molecular nodes in glycan-mediated immune synapses, foreseeing their application in deciphering tumor-immune interactions.

## Introduction

The ability of cancer cells to evade immune surveillance is frequently mediated by aberrant glycosylation, such as the overexpression of truncated *O*-glycans like the Tn antigen (GalNAcα1-*O*-Ser/Thr). These alterations enable tumor cells to exploit immune checkpoints, disrupt antigen presentation, and reprogram antigen-presenting cells (APCs) towards more cancer-tolerogenic and immunosuppressive phenotypes (1). This glycan-mediated immune evasion contributes to tumor progression and resistance to immunotherapy, highlighting the importance of understanding and targeting these glycosylation patterns and their associated receptors in cancer treatment strategies. However, the glycoproteins mediating this immunosuppression, their cognate immune cell receptors, and the resulting downstream synaptic events remain poorly characterized, hampering a broader understanding of cancer-immune cell crosstalk and innovative immune checkpoint inhibition strategies. Here, we introduce GlycoAvatars, an innovative platform for studying interactions between cancer glycoproteomes and immune receptors, providing a decisive tool to advance the identification of key molecular players in glycan-mediated immune evasion and to uncover new targets for immunotherapy.

## Results and Discussion

We developed GlycoAvatars—magnetic beads coated with plasma membrane glycoproteins from cancer cells—to investigate glycan-mediated interactions with immune cells under physiological conditions (**Fig. 1 A**). This approach preserves native receptor conformation and co-receptor associations. To model the effects of altered glycosylation in aggressive cancers, we generated GlycoAvatars using Tn-expressing glycoproteins from *C1GALT1* knockout (KO) AGS (gastric cancer) and SW480 (colorectal cancer) cells, which exhibit homogenous Tn antigen expression in contrast to the extended glycosylation found in wild-type (WT) cells (**Fig. 1 B-D**). Tn-expressing glycoproteins were isolated from plasma membrane-enriched extracts using VVA lectin affinity after PNGase-F digestion to remove N-glycans, to mitigate the influence of other major types of protein glycosylation. Immature and mature monocyte-derived dendritic cells (iDCs and mDCs) and immature macrophages (iMACs) were exposed to these GlycoAvatars (**Fig. 1 E**). The cell-bead conjugates were treated with a membrane-impermeable crosslinker to stabilise the receptor-ligand-interactions before cell lysis and proceeding. Proteins interacting with the GlycoAvatars were identified using a membrane-impermeable crosslinker (BS3), magnetic pulldown, and nanoLC-MS/MS (2).

**Figure 1.**
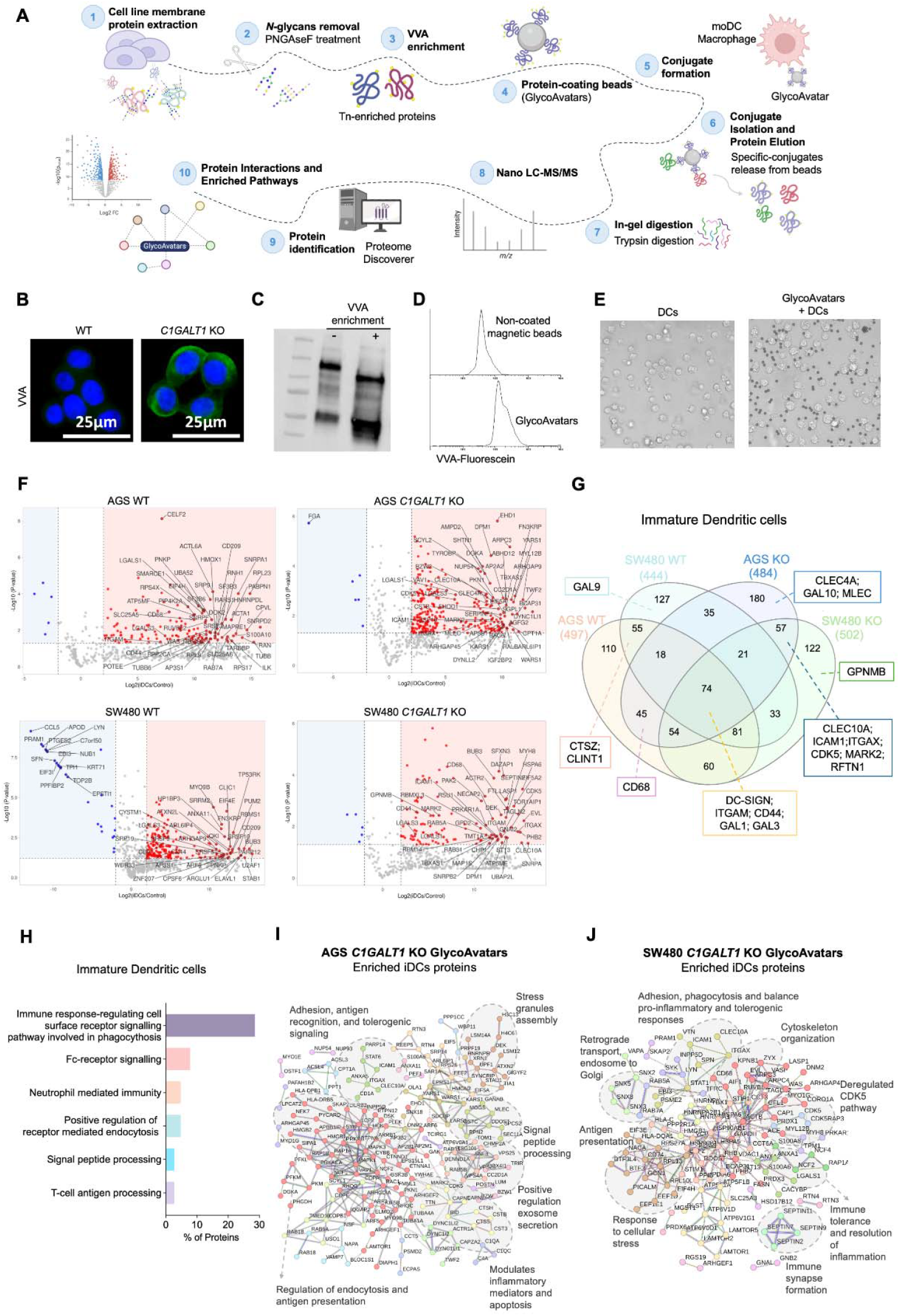
Mass spectrometry roadmap for identifying glycoprotein-mediated synapses between immune cells and engineered GlycoAvatars. **A. Workflow overview for immune cell receptor identification**. Plasma membrane protein extracts (1) derived from glycoengineered cancer cell lines (AGS and SW480 C1GALT1KO) reflecting a cancer-associated *O*-glycome were *N*-deglycosylated (2) and enriched using VVA lectin affinity chromatography (3) to isolate Tn-enriched glycoproteins. Wild-type (WT) extracts plasma membrane proteins were only *N*-deglycosylated without considering the enrichment step. The enriched proteins were conjugated to magnetic beads to create GlycoAvatars (4). GlycoAvatar formation was validated using flow cytometry. Subsequently, GlycoAvatars were incubated with monocyte-derived dendritic cells (moDCs) or macrophages (MACs) (5) and after confirming specific binding between immune cells and GlycoAvatars, unbound cells were removed, immune cell membranes disrupted, and after washing, the protein-protein conjugates were eluted from the beads (6). The eluates underwent in-gel trypsin digestion (7), mass spectrometry acquisition (8), and bioinformatic analysis, including protein identification (9), comprehensive protein-protein networks and biological process pathways (10). **B. Glycoengineered cancer cell lines**. AGS and SW480 cell lines, engineered to overexpress the cancer-associated Tn antigen through *C1GALT1* knockout (inhibiting core 1 *O*-glycan synthesis and subsequent chain elongation), served as cancer models. **C. Tn-glycoprotein enrichment**. Plasma membrane protein extracts from the *C1GALT1* KO models were enriched by VVA-agarose lectin affinity to specifically isolate glycoproteins bearing the Tn antigen. **D. GlycoAvatars validation**. Successful glycoprotein coating and Tn antigen availability on the GlycoAvatars were confirmed by flow cytometry using VVA-Fluorescein lectin staining. **E. Formation of immune synapses**. Interaction and conjugate formation between DCs or MACs and GlycoAvatars were validated via inverted bright-field microscopy. **F. Identification of specific synaptic proteins**. Employing Volcano plots, the significantly enriched proteins in iDC-GlycoAvatar (WT or *C1GALT1* KO) interactions were identified as synapse-specific. The robustness of this method was demonstrated by consistently detecting CLEC10A (MGL) in DC-GlycoAvatars conjugates from AGS and SW480 *C1GALT1* KO models across donors (n=3). In contrast, CLEC10A was absent in conjugates formed with WT GlycoAvatars, corroborating specificity. **G. Venn diagram depicting protein distribution from GlycoAvatar-immature dendritic cell interactions**. Proteins enriched in each condition were selected based on statistical significance (Log_2_ fold change (iDCs / Controls) > 2 and -Log_10_ p-value > 1.3). **H. Functional enrichment of iDC-specific proteins**. Enriched pathways and biological processes from DC-specific proteins interacting with KO GlycoAvatars. **Protein-protein interaction network of iDC-specific proteins from AGS (I) and SW480 (J) *C1GALT1* KO GlycoAvatars**.

GlycoAvatar pulldowns from AGS and SW480 *C1GALT1* KO cell lines identified 237 and 179 proteins in iDCs, respectively, that were significantly enriched or exclusively expressed compared to cells with extended glycosylation (**Fig. 1 F-G and Table S1**). Approximately 60% of these proteins were predicted to be plasma membrane-associated, expanding our understanding of immune cell molecules potentially interacting with Tn-expressing glycoproteins GlycoAvatars’ exclusive proteins included CLEC10A (CD301 or MGL) and CLEC4A (DCIR), which are well-known Tn-binding lectins mediating immune suppression (3) (**Fig 1 F-G**), thus validating the approach. Other potential candidates with relevant reported roles in DCs functions include ICAM1 (CD54) (4) and ITGAX (CD11c) (5) (involved in synapse formation, phagocytosis, and migration), MARK2 (PAR-1b/EMK) (6, 7) (regulates pro-inflammatory cytokine responses), and RFTN1 (Raftlin) (8) (facilitating Toll-like receptor trafficking and interferon production). Furthermore, GAL10 (AGS KO) and GPNMB (SW480 KO) were identified and are known to promote immune tolerance and inhibit DC maturation (9, 10). We also observed other immune lectins, including DC-SIGN (CD209), MLEC, several galectins (GAL1, GAL3), and CD44 (**Fig. 1 F-G**). The interaction of these glycoproteins (some also detected in WT avatars) with the Tn antigen warrants further investigation. Pathway analysis revealed significant enrichment for biological processes related to receptor activation and antigen uptake (phagocytosis, Fc-receptor signaling, receptor-mediated endocytosis) (**Fig. 1 H**). In addition, we identified a variety of intracellular proteins (around 40% of the interactome) likely involved in the immune synaptic cascade triggered by Tn-expressing glycoproteins. The consistent identification of the intracellular protein CDK5 a is particularly noteworthy, given its established role in regulating DC migration, antigen presentation, and immune responses via its influence on cytoskeletal dynamics and endocytosis (11). Furthermore, the identification of numerous intracellular proteins alongside membrane receptors strongly suggests a functional, rather than incidental, involvement in glycan-mediated immune signaling. This conclusion is supported by STRING analysis revealing significant connectivity between the identified membrane and intracellular proteins (**Fig. 1 I-J**), indicative of an integrated immune synaptic cascade. It is also reinforced by the cell-line-specific interactomes (**Fig. 1 I-J**), highlighting the context-dependent nature of these interactions. In summary, the GlycoAvatar platform identified proteins potentially involved in glycan-driven interactions between iDCs and aggressive cancer cells. Future studies will be necessary to determine their specific roles and functional significance.

We further extended our approach to investigate the impact of cell maturation state and cell type by comparing immature and mature DCs and immature MACs. Distinct glyco-interactomes were observed between iDCs and mDCs for both cell lines (**Fig. 2 A**), with CLEC10A and CLEC4A showing significantly higher enrichment in iDCs, consistent with their developmental downregulation. This observation underscores the importance of considering the maturation state when investigating receptor-ligand interactions and highlights the sensitivity of the GlycoAvatar platform in capturing these nuances. Both cell lines showed enrichment in negative regulation of interferon type I signaling pathways (**Fig. 2 B**), suggesting immune suppression mediated by the Tn antigen. However, cell-type-specific differences emerged. AGS cells showed enrichment in IL-10 production and Th1 T helper (Th) responses, while SW480 cells showed enrichment in B-cell differentiation pathways (**Fig. 2 B**). These findings support the known immunosuppressive role of the Tn antigen and suggest cell-specific mechanisms underlying this effect. Despite these cell-type-specific differences, several proteins consistently mediating key immune processes related to tumor dynamics were also identified across both cell types (**Fig. 2 C**). These included ICAM1 (immune activation), EBI3 (immunosuppression), OAS3 (interferon responses), and proteins associated with DC migration and immune synapse formation (MARCKS, WIPF1) (12-14). Many of these interactions are novel in the context of Tn antigen recognition. This interplay appears to balance immune activation and suppression, warranting further investigation into the context-dependent mechanisms. In iMACs, we identified 73 and 50 proteins from AGS and SW480-derived GlycoAvatars, respectively (**Table S2**). Notably, only a small subset of membrane proteins was common to iDCs, including MARK2, ITGAX, and GPNMB as well as CDK5 (**Fig. 2 D-E**), further supporting their role in Tn antigen recognition and APC modulation. Key iMAC-enriched cytoplasmic proteins further included RHOC and PSTPIP2. STRING analysis (**Fig. 2 F**) revealed their involvement in cytoskeletal rearrangement and antigen uptake (CFL1, ITGAX, ANAX5, RHOC) and inflammatory responses modulation (CDK5, CAPG, GPNMB, MARK2, PSTPIP2) (15). These proteins were also implicated in pathways associated with M1/M2 polarization (GPNMB, MARK2), pro-tumoral behavior (RHOC, CDK5, PIK3C2A), and anti-inflammatory responses (PHB) (16), along with antigen presentation (HLA-DRA, HLA-DRB1) and cell survival/adhesion. Furthermore, main biological functions revealed roles in neutrophil activation, Fc-gamma receptor signaling, immunological synapse formation, and macrophage activation (**Fig. 2 G**). These findings align with the pro-tumoral role of macrophages in Tn-expressing cancers and suggest potential therapeutic targets. The distinct pathway enrichments in iMACs versus iDCs further highlight the GlycoAvatar platform’s capacity to reveal context-specific immune interactions.

**Figure 2.**
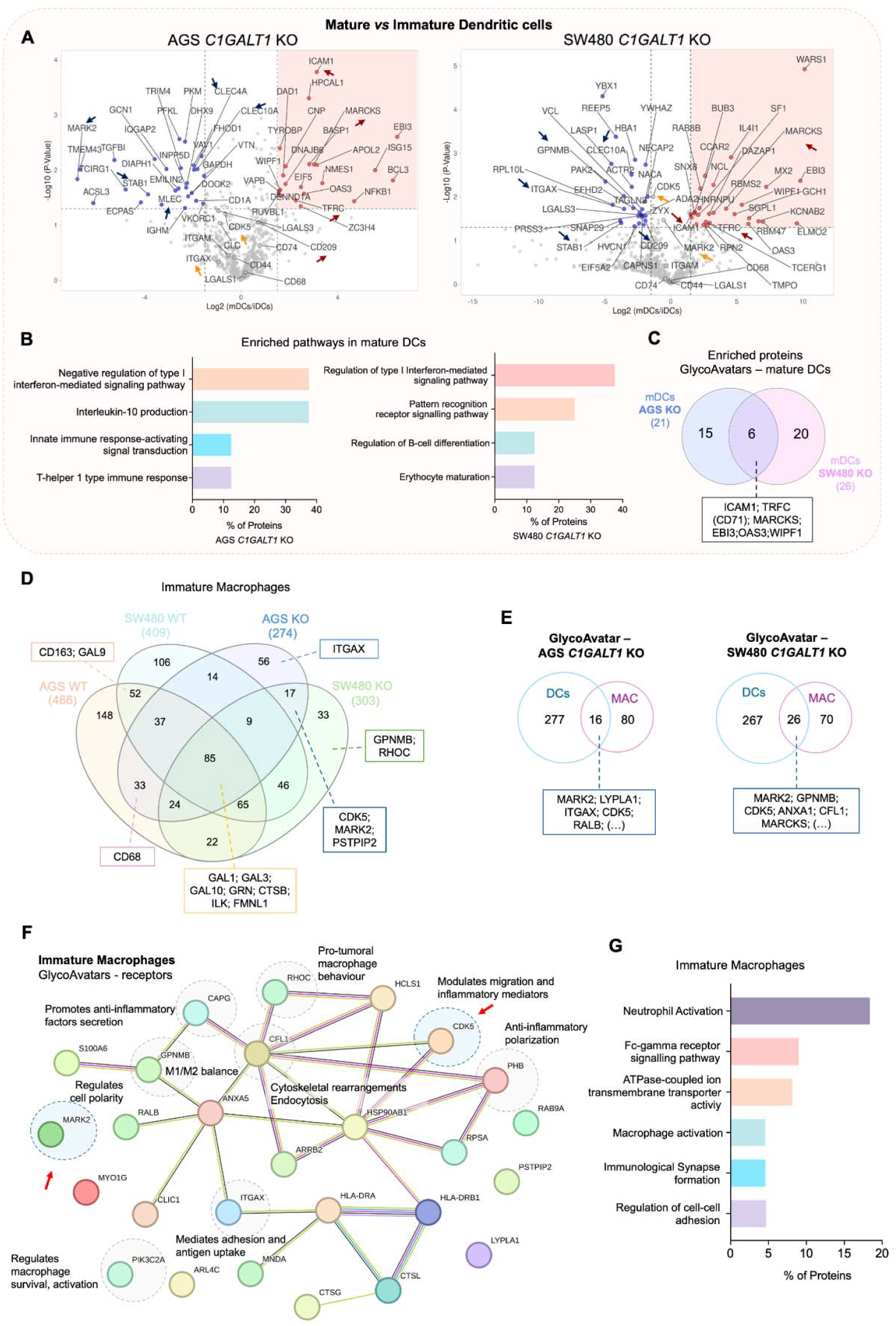
Identification and characterization of proteins mediating GlycoAvatar-immune cell synapses using mass spectrometry. **A. Differential protein expression between DC maturation states**. Comparative analysis of protein abundances in mature DCs (mDCs) versus immature DCs (iDCs) revealed maturation state-specific proteins. For instance, CLEC10A was more abundant in iDC-GlycoAvatar (*C1GALT1* KO) conjugates than in mDC-GlycoAvatar conjugates, consistent with literature reports that CLEC10A expression is higher in iDCs. Blue arrows: indicate some relevant downregulated proteins, orange arrows: non-significantly changed and red arrows: upregulated proteins in mDCs. **B. Enriched pathways in mDC interactions**. Proteins significantly enriched in mDC interactions with AGS and SW480 *C1GALT1* KO GlycoAvatars were associated with biological pathways such as negative regulation of type I interferon signalling, innate immune signalling, and other immune regulatory processes. **C. Common proteins across cell lines**. Among the proteins enriched in mDC interactions with AGS (21 proteins) and SW480 (26 proteins) *C1GALT1* KO GlycoAvatars, 6 proteins were common, highlighting shared mechanisms in the Tn-specific synapses. **D. Venn diagram depicting protein distribution from GlycoAvatar-immature macrophage interactions**. Proteins exclusive to AGS/SW480 *C1GALT1* KO GlycoAvatars-immature macrophages conjugates highlight specific interactions driven by Tn-enriched glycoproteins, whereas those exclusive to WT GlycoAvatars conjugates reflect interactions associated with the native *O*-glycome. **E. Summary of DC- and MAC-specific proteins from KO-GlycoAvatars**. Venn diagrams highlight the overlap and exclusivity of proteins identified in iDCs and iMAC interacting with AGS and SW480 KO GlycoAvatars. **F. Protein-protein interaction network of MAC-specific proteins from KO GlycoAvatars**. STRING-based analysis of MAC-specific proteins revealed key regulators of macrophage polarization (RHOC, MARK2), migration (PSTPIP2, CDK5), and cytoskeletal dynamics (CFL1, ITGAX). **G. Functional enrichment of MAC-specific proteins**. Enriched pathways and biological processes from MAC-specific proteins interacting with KO GlycoAvatars.

### Concluding Remarks

This study establishes the GlycoAvatar platform as a powerful tool for characterizing glycan-mediated immune interactions in cancer under physiological conditions. Our approach, employing magnetic beads coated with cancer cell plasma membrane glycoproteins, maintains native receptor conformations and co-receptor associations, offering a more biologically relevant model than traditional methods. The use of glycoproteins from *C1GALT1* knockout cancer cells allowed us to specifically investigate the role of Tn-carrying glycoproteins in this interaction, which is a critical challenge in cancer therapy development. This study reveals novel, proteome-dependent interplays between glycosylation and immune cell function, as demonstrated by the identification of 495 proteins (223 membrane-associated) potentially interacting with immaturely glycosylated glycoproteins on cancer cells, influencing immune synapse formation. Analysis revealed both common and cell-type-specific protein networks significantly modulated by the Tn antigen, greatly expanding our understanding of glycan-mediated immune modulation. The crucial role of cellular context (cell type and maturation state) in these interactions is highlighted by these findings, suggesting potential therapeutic targets for innovative immunotherapies, offering new insights into cancer-immune cell crosstalk. The identification of key proteins involved in immune activation (ICAM1; RHOC), suppression (EBI3; GPNMB; PHB), and interferon-stimulated responses (OAS3; RFNT1) highlight potential therapeutic targets. The consistent presence of several proteins influencing DC migration and immune synapse formation (MARCKS, WIPF1) underscores the multifaceted role of Tn-expressing glycoproteins in shaping the immune response. Furthermore, the significant involvement of CDK5, a key regulator of dendritic cell migration and antigen presentation, reveals the potential for intracellular signaling pathways to mediate glycan-driven immune interactions. Collectively, we suggest relevant protein nodes within these networks holding potential for precise modulation of immune responses in Tn-expressing cancers. For instance, the observed cell-line-specific interactomes highlight the potential for personalized therapies tailored to unique glycoproteoforms-mediated interactions. This also underscores the importance of considering cell-type-specific responses and the nature of the glycoproteome rather than just the nature of the glycan chain in designing effective immunotherapies. Future research should focus on: (1) validating Tn-lectin binding; (2) analyzing post-translational modifications beyond *O*-glycosylation; and (3) exploring the therapeutic potential of the identified proteins. The GlycoAvatar platform thus offers a relevant approach for identifying novel immune checkpoints and developing targeted cancer therapies.

## Materials and Methods

### Glycoproteins Isolation

Plasma membrane proteins were extracted, quantified, and adjusted to 300 μL with TBS-0.3% SDS and 15 μL of 20% NP-40. Proteins were deglycosylated overnight at 37 °C with PNGase F (2 U/20 μg protein), followed by enzyme inactivation at 95 °C for 20 minutes. Samples were diluted to 400 μL with LacA buffer (20 mM Tris, pH 7.4, 150 mM NaCl, 1 mM Urea, 0.1 M CaCl_2_, 0.1 M MgCl_2_, 0.1 M MnCl_2_, 0.1 M ZnCl_2_) incubated with 100 μL VVA-agarose for 30 minutes, then washed and eluted with 3% acetic acid. Eluates were dried using a SpeedVac and reconstituted in TBS-0.3% SDS.

### GlycoAvatars development

Dynabeads™ M-450 Tosyl-activated (20×10□) were washed twice with Buffer 1 (0.1 M phosphate buffer, pH 7.4–8.0) and incubated with 10 μg of target membrane proteins for 1.5 hours at room temperature with agitation (1000 rpm). Buffer 1 containing 1% BSA (final 0.167% BSA) was added, and the beads were incubated overnight at 37 °C to ensure binding. Beads were washed with 20 mM HEPES and incubated for 2 hours at room temperature in 25 mM (bis(sulfosuccinimidyl)suberate) BS3 crosslinker, then washed with Buffer 3 (0.2 M Tris, 0.1% BSA, pH 8.5) and incubated overnight to deactivate residual tosyl groups. After final washes with Buffer 2 (PBS, 0.1% BSA, 2 mM EDTA, pH 7.4), binding of Tn-enriched proteins was confirmed by flow cytometry using VVA. Beads were stored in Buffer 2 at 4 °C for up to one month.

### GlycoAvatars-Immune Cells Complexes and Immune Synapse isolation

Dendritic cells or macrophages were resuspended in RPMI medium without phenol red and combined with protein-coated beads at a 1:1 ratio, incubating for 60 minutes at 37°C with agitation (1000 rpm). Conjugation was confirmed microscopically, and the conjugates were washed with cold CSK buffer (300 mM sucrose, 100 mM sodium chloride, 10 mM PIPES, pH 6.8, 3 mM magnesium chloride) to remove unbound cells. After resuspension in CSK buffer with 0.5% Triton X-100 and protease and phosphatase inhibitors, the conjugates were sonicated and repeatedly washed with CSK buffer 5% Triton X-100 to remove debris. Bound proteins were eluted with 25 μL of Laemmli Buffer at 70°C for 30 minutes (1000 rpm). The eluates were stored at -20 °C for up to one week.

### Proteomics

Eluates from GlycoAvatars-Immune cells protein complexes were identified by a bottom-up proteomics strategy by nanoLC-HCD-M/MS. Briefly, isolated proteins were loaded into electrophoresis gels and stained using a Zinc Reversible Stain Kit. Gel bands were excised into 1–2 mm pieces, dissolved in Tris-glycine buffer pH 8.0, and washed thrice with ultrapure water for 10 minutes each. The gel pieces were dehydrated with LC-MS-grade acetonitrile (ACN) for 5–10 minutes. Disulfide bonds were reduced with 20 mM dithiothreitol (DTT) at 56 °C for 30 minutes, followed by dehydration with 100% ACN. Rehydration with 55 mM iodoacetamide (IAA) for 20 minutes in the dark prevented bond reoxidation. After washing with 100 mM ammonium bicarbonate and further dehydration, enzymatic digestion was initiated with 0.02 μg/μL trypsin in 40 mM ammonium bicarbonate/10% ACN at 37 °C overnight in a humid chamber, followed by enzyme inactivation in 100% ACN. Peptides were extracted with 50% ACN/5% formic acid (FA) two times, and the extracts were dried using a SpeedVac.

### Mass spectrometry

For protein analysis, peptides were loaded into a Vanquish Neo UHPLC system coupled with a QExactive Plus Hybrid Quadrupole-Orbitrap mass spectrometer (Thermo Fisher Scientific), which was fitted with a nano-electrospray ion source (EASY-Spray source; Thermo Fisher Scientific). Mobile phases were 0.1% FA in ultrapure water (eluent A) and 0.1% FA in 80% ACN (eluent B). A 10 μL sample was injected into a trapping column (C18 PepMap Neo, 5 μm particle size 300μm × 5mm) and separated on an analytical column (EASY-Spray C18 PepMap, 100 Å, 75 μm × 150 mm, 3 μm particle size) at 0.25 μL/min and 35 °C. Peptide separation used a gradient of 2.5%– 12% B over 7 min, 12%–46% B over 50 min, and 46%–99% B over 5 min, holding at 99% B for 10 min. Mass spectrometry operated in positive ion mode (m/z 300–2000) with a 1.9 kV spray voltage and 275 °C capillary temperature. Full MS settings included a 140,000 resolution, AGC target of 3×10□, and 200 ms maximum injection time, with the 15 most intense ions selected for higher energy collisional dissociation (HCD) with NCE of 30%. Data were recorded with Xcalibur (v4.5).

### Protein Annotation and Data Analysis

Data were analysed using the Proteome Discoverer (v3.1) with SequestHT search engine and Percolator against the SwissProt human proteome database (accessed on October 22nd, 2023). Parameters included trypsin specificity, up to two missed cleavages, a 10-ppm precursor ion tolerance, a 0.02 Da product ion tolerance, fixed carbamidomethylcysteine (+57.021 Da), and variable methionine oxidation (+15.995 Da). Proteins with fewer than two unique peptides or low/medium FDR confidence were excluded to generate the final curated protein list. Proteins were annotated based on gene ontology (GO) terms for cellular location, molecular functions, and biological processes using Proteome Discoverer 3.1. Proteins identified in the eluates from conditions containing only GlycoAvatars (AGS/SW480 WT and *C1GALT1* KO) were used as controls to profile proteins from tumor cell lines and evaluate their relative abundance. Significantly enriched proteins from DC/Mac-GlycoAvatar interactions were identified using Volcano Plots (VolcaNoseR), selecting proteins with a Log2 fold change > 2 (Interactions/Coated Beads) and an adjusted p-value < 0.05 (-Log10 p-value > 1.3). Proteins uniquely found in DC/Mac-AGS/SW480 *C1GALT1* KO and not present in the AGS/SW480 WT interactions samples were further analysed for their protein-protein interactions, molecular functions, biological processes, and KEGG pathways using STRING version 12.0. A network of biological processes for these proteins was created using the ClueGO plugin in Cytoscape (v3.10.2), considering significant pathways (p ≤ 0.05), a GO tree interval of 4–10, and a kappa score of 0.64 for pathway clustering.

## Acknowledgments

The authors wish to acknowledge the Portuguese Foundation for Science and Technology (FCT) for 2022.12980.BD (AM), SFRH/BD/146500/2019 (DOI 10.54499/SFRH/BD/146500/2019) and COVID/BD/153652/2024 (MRS), and Principal researcher contract 2022.08311.CEECIND (JAF). FCT is co-financed by European Social Fund under Human Potential Operation Programme from National Strategic Reference Framework. The authors also acknowledge FCT/MCTES funding within the projects RESOLVE (https://doi.org/10.54499/PTDC/MED-OUT/2512/2021) and funding for the IPO research center (PEst-OE/SAU/UI0776/201) as well as institutional funding (CI-IPOP-29-2016-2022, CI-IPOP-58-2016-2022). This article is also supported by project NORTE-01-0145-FEDER-000012 and “The Porto Comprehensive Cancer Center Raquel Seruca” with the reference NORTE-01-0145-FEDER-072678 - Consórcio PORTO.CCC—Porto.Comprehensive Cancer Center Raquel Seruca, under the PORTUGAL 2020 Partnership Agreement, through the European Regional Development Fund. We further acknowledge the Research Council of Finland (InFLAMES flagship, decision numbers: 337530 and 357910; and project funding to PKM, decision number: 339810) and Finnish Cultural Foundations (to DMC).

